# Interaction between neuronal encoding and population dynamics during categorization task switching in parietal cortex

**DOI:** 10.1101/2020.08.20.259820

**Authors:** Krithika Mohan, Oliver Zhu, David Freedman

## Abstract

Primates excel at categorization, a cognitive process for assigning stimuli into behaviorally relevant groups. Categories are encoded in multiple brain areas and tasks, yet it remains unclear how neural encoding and dynamics support cognitive tasks with different demands. We recorded from parietal cortex during flexible switching between categorization tasks with distinct cognitive and motor demands, and also studied recurrent neural networks (RNNs) trained on the same tasks. In the one-interval categorization task (OIC), monkeys rapidly reported their decisions with a saccade. In the delayed match-to-category (DMC) task, monkeys decided whether sequentially presented stimuli were categorical matches. Neuronal category encoding generalized across tasks, but categorical encoding was more binary-like in the DMC task and more graded in the OIC task. Furthermore, analysis of the trained RNNs supports the hypothesis that binary-like encoding in the DMC task arises through compression of graded feature encoding by population attractor dynamics underlying short-term working memory.

## Introduction

Visual categorization is a cognitive process in which continuous visual stimuli are associated with discrete groups according to their behavioral relevance. Visual categories are encoded in multiple brain regions in the primate, including the posterior parietal cortex (PPC) (Freedman and Assad, 2006), prefrontal cortex (PFC) (Freedman et al., 2001; Swaminathan and Freedman, 2012), and inferotemporal cortex (ITC) (Freedman et al., 2003; Meyers et al., 2008) with single neuron activity showing distinct responses to stimuli in different categories. Yet, the format of neural category representations differs between these brain areas. Whereas PPC and PFC encode categories in an abstract or binary-like format with nearly identical firing rates to all stimuli within a category (Freedman et al., 2006; Swaminathan and Freedman, 2012), ITC neurons encode categories with more graded firing rates to stimuli within a category, thus ‘mixing’ abstract category signals with sensory feature encoding (Freedman et al., 2003; Meyers et al., 2008). These two formats of category encoding – binary or graded – reflect a tradeoff between sensory feature encoding and abstract category encoding, yet the mechanistic origins of this tradeoff remain unknown.

Category signals have been studied through behavioral tasks that divide a continuous range of sensory stimuli into two or more discrete categories by an arbitrary category boundary (Freedman et al., 2001; Freedman and Assad, 2006). Further, these tasks decouple categorical decisions from animals’ motor responses by using delayed matching paradigms which require comparing sequentially presented stimuli separated by a delay (Freedman et al., 2001; Freedman and Assad, 2006). For example, in the delayed match-to-category (DMC) task, monkeys must release a manual touch-bar if a test stimulus matches the category of a previously presented sample. During DMC tasks, neurons in PFC and lateral intraparietal area (LIP) of PPC (Freedman and Assad, 2006; Swaminathan and Freedman, 2012), show binary-like categorical encoding of motion-dot stimuli, and LIP activity in particular correlates with animals’ categorical choices on a trial-by-trial basis (Swaminathan and Freedman, 2012).

Recent work using reversible cortical inactivation demonstrated a causal role for LIP in visual categorical decisions (Zhou and Freedman, 2019). These binary-like category signals have mostly been observed in tasks with a short-term memory-delay period which helps dissociate sensory signals from decision and motor signals. However, the delay also imposes additional mnemonic demands, that are not directly related to categorization per se. Thus, it remains unknown if these binary-like category signals evident in LIP emerge because of the cognitive demands of delayed matching paradigms (e.g. short-term memory or matching) predominantly used in previous categorization studies. This raises two related questions – 1) does LIP’s role in categorical decisions generalize across tasks with different task demands and 2) does the format of category encoding depend on changing task demands?

Here, we examine whether LIP plays a generalized role in categorical decisions across different categorization task paradigms, and whether categorical encoding is modulated by task demands. We devised a cued task switching paradigm in which monkeys alternated in blocks between two motion categorization tasks that varied in their task demands. In both tasks, monkeys grouped 360° of motion directions into two categories according to the same arbitrary, learned category boundary. The first task was a one-interval categorization (OIC) task, in which monkeys rapidly reported their categorical decisions with a saccade to a red (category one) or green (category two) target. The second task was a DMC task which required monkeys to compare sequentially presented sample and test stimuli that were separated by a delay, and report whether they matched in category with a manual response. The tasks varied in: - (i) the report of direct (OIC) versus sequential (DMC) category judgments, (ii) associative mappings between the stimulus category and saccade target colors (OIC) and working memory and sequential comparison (DMC) and (iii) the effector for the motor response of eye (OIC) versus hand movements (DMC).

Single neurons and populations in LIP showed strong and similar category encoding in both tasks, indicating that LIP plays a generalized role in categorical decision-making. However, the format of categorical encoding was more abstract with binary-like neural activity in the DMC task but more mixed with graded neural activity in the OIC task. Furthermore, categorical encoding was more temporally stable and sustained in the DMC task. We hypothesize that this binary-like format of category encoding in the DMC task is generated by population attractor dynamics required to maintain stimulus information over memory delay periods. Because only the DMC task requires maintaining category information in working memory, attractor dynamics appears to compress category-related information to a simpler, binary format by collapsing all directions within a category towards a single population state. We validated this hypothesis using recurrent neural networks trained to perform both tasks and found that analysis of the fixed-point structure of these RNNs revealed that following stimulus onset, sample categories produce stable attractors in the DMC, but not in the rapid OIC task. Thus, the binary-like categorical responses observed in DMC tasks may not be necessary for categorization per se, but may instead arise from the network dynamics underlying the storage of stimulus information in working memory. More broadly, our approach of incorporating flexibility in behavioral tasks facilitates the understanding of how neural activity patterns support diverse cognitive computations and distinguishes between task-specific encoding underlying categorization and working memory.

## Results

### Categorization task switching behavior

Two monkeys alternated between the OIC and DMC tasks in which they categorized the same random-dot motion stimuli in blocks of 20 or 30 correct trials in monkey B or M, respectively (Figure 1a). In both tasks, 360° of motion directions were divided into two categories based on an arbitrary learned category boundary, with 5 motion directions in each category (Figure 1b). On every trial, after a 500 ms fixation period, one sample stimulus was presented for 500 ms in the OIC task, and for 650 ms in the DMC task in the neuron’s receptive field (RF). Both tasks were visually identical until 500-ms from sample onset (the shared sample period) after which they diverged. In the OIC task, saccade targets (red and green squares) appeared at 500 ms after sample onset and monkeys reported their categorical decisions with a saccade to either the red or the green target to indicate category one or two, respectively. Importantly, the target position varied randomly on every trial between two possible locations – one inside the RF and the other diametrically opposed to the RF. This ensured a dissociation between the categorical decision (red versus green) and the motor responses (saccade towards versus away from the RF). In the DMC task, the sample stimulus was followed by a one second delay and a test stimulus, and monkeys reported whether the test stimulus was a categorical mech to the previously presented sample stimulus by releasing a touch bar. Each test stimulus was randomly sampled from among the ten possible motion directions (five in each category).

**Figure 1.**
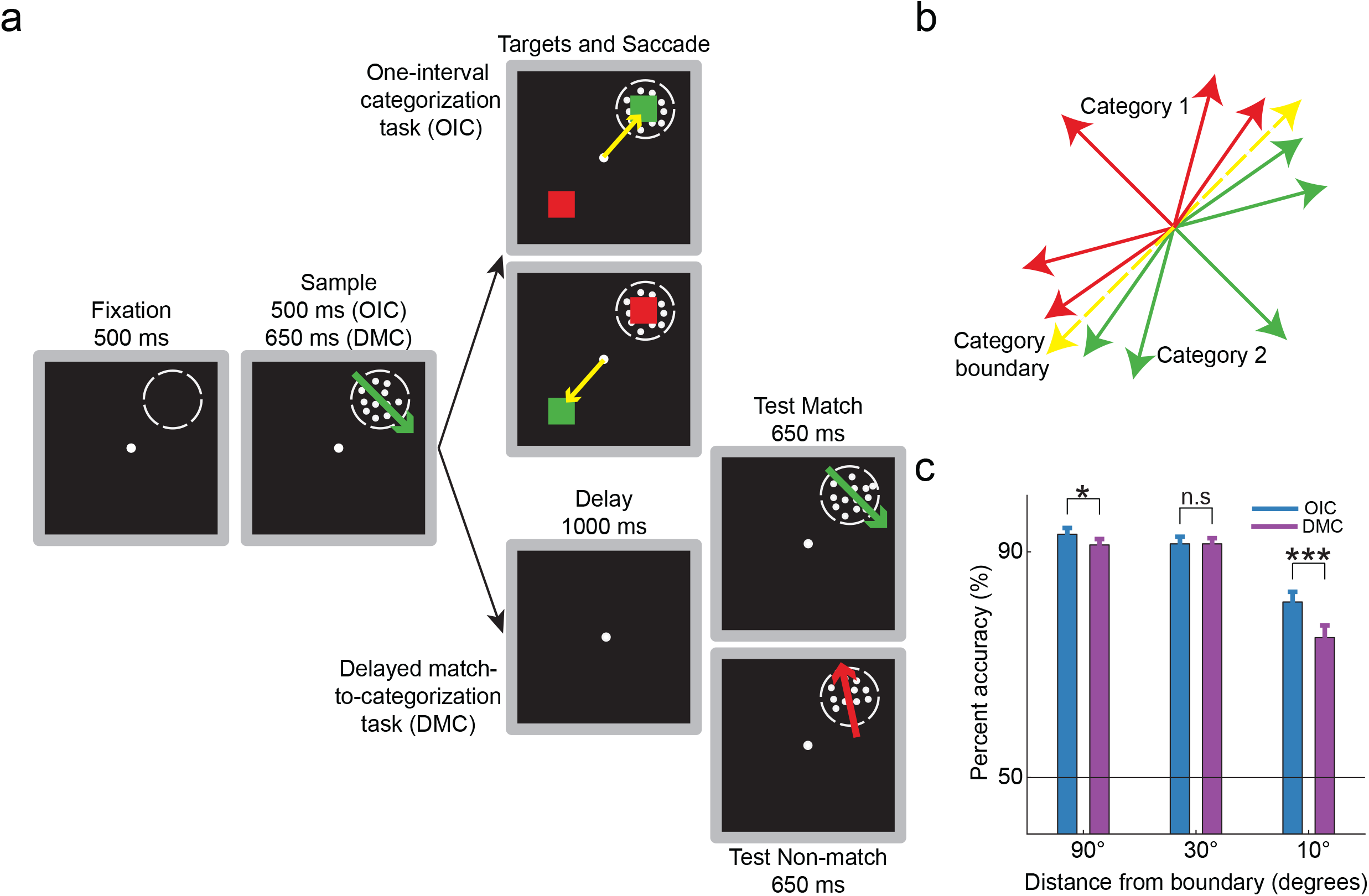
Categorization task switching and behavior. (a) Monkeys alternated between the one-interval categorization (OIC) task and the delayed match-to-category task in blocks of 20 trials. On each trial, a 500 ms fixation period was followed by a sample period (500 ms in OIC, 650 ms in DMC) during which moving dots appeared in the receptive field (RF) of the recorded neuron. In OIC, colored saccade targets (red and green –see 1b) previously associated with the two motion categories appeared at 500 ms from sample onset. Once the targets appeared, monkeys made a saccade to the red or green target to indicate category one or two, respectively. In DMC, the sample stimulus was followed by a one second delay, and one or two test stimuli. If the sample and test stimulus matched in category, monkeys released a touchbar, otherwise monkeys held the touchbar until a second test stimulus appeared, which was always a categorical match. (b) The stimuli were random dots moving in one of ten possible motion directions, divided into two categories (represented by red and green colors) by an arbitrary category boundary (yellow dashed line). (c) Monkeys performed categorization task switching successfully at 87.6% correct in OIC, 84.13% correct in DMC. Performance increased with distance from the category boundary in both tasks. Performance in OIC was significantly greater than DMC for the easiest (90° away from boundary) and hardest (10° away from boundary) motion directions. Error bars indicate s.e.m. * p<0.05, *** p<0.001, n.s not significant, unpaired two-sample t-test.

Based on a colored cue presented at the start of each trial, monkeys switched between the OIC and DMC tasks in blocks. Monkeys successfully switched between the two tasks with a mean behavioral performance of 87.6% correct in OIC (Monkey B: 86.4%; Monkey M: 88.9%) and 84.3% correct in DMC (Monkey B: 83.1%; Monkey M: 85.7%) (Figure 1c). Performance of both monkeys was significantly higher in the OIC task for the near-boundary directions that were 10° away from the boundary (p-value of paired t-test: 10^−08^).

### LIP neurons are preferentially engaged in the OIC task

We recorded from 100 LIP neurons (monkey B, N=44 single units; monkey M, N=56 single units) as monkeys alternated between the OIC and DMC tasks. We compared LIP responses in the OIC and DMC tasks to determine whether LIP neurons were preferentially engaged in one task over the other. We hypothesized that LIP’s well-known role in spatial attention and saccadic eye movements would be reflected in the firing rates in the OIC task, which requires a saccadic report (Gnadt and Andersen, 1988; Colby, Duhamel and Goldberg, 1996; Bisley and Goldberg, 2010). On the other hand, LIP’s well-known role in spatial functions might be distinct from its role in visual motion categorization, as suggested by a previous study in which spatial signals and category signals independently influenced LIP activity, when the saccadic report and the categorical report were behaviorally not linked (Rishel, Huang and Freedman, 2013). To compare firing rates between the OIC and DMC tasks, we focused on neural responses during the first 500 ms of the sample period during which monkeys were fixating and stimulus presentation was identical, henceforth referred to as the shared sample period. Out of 100 recorded neurons, 82 neurons showed significant differences in firing rates between the OIC and DMC tasks in the shared sample period. Among the 82 neurons that showed significant firing rate differences between the two tasks, 73.2% (60/82) of individual LIP neurons responded with higher mean firing rates in the OIC task (unpaired two sample t-test, p<0.01).

Further, firing rates of individual neurons in the OIC task were positively correlated with firing rates in the DMC task (Figure 2c, 2d, r^2^ = 0.95, p<10^−4^) and these values were well fit by a linear model (R^2^ = 0.90) with positive slope 1.13 (95% confidence intervals: (1.05,1.22)) and intercept 2.7 (95% confidence intervals: (−0.56,6.06)), indicating that OIC firing rates were significantly higher than DMC firing rates in the LIP population.

**Figure 2.**
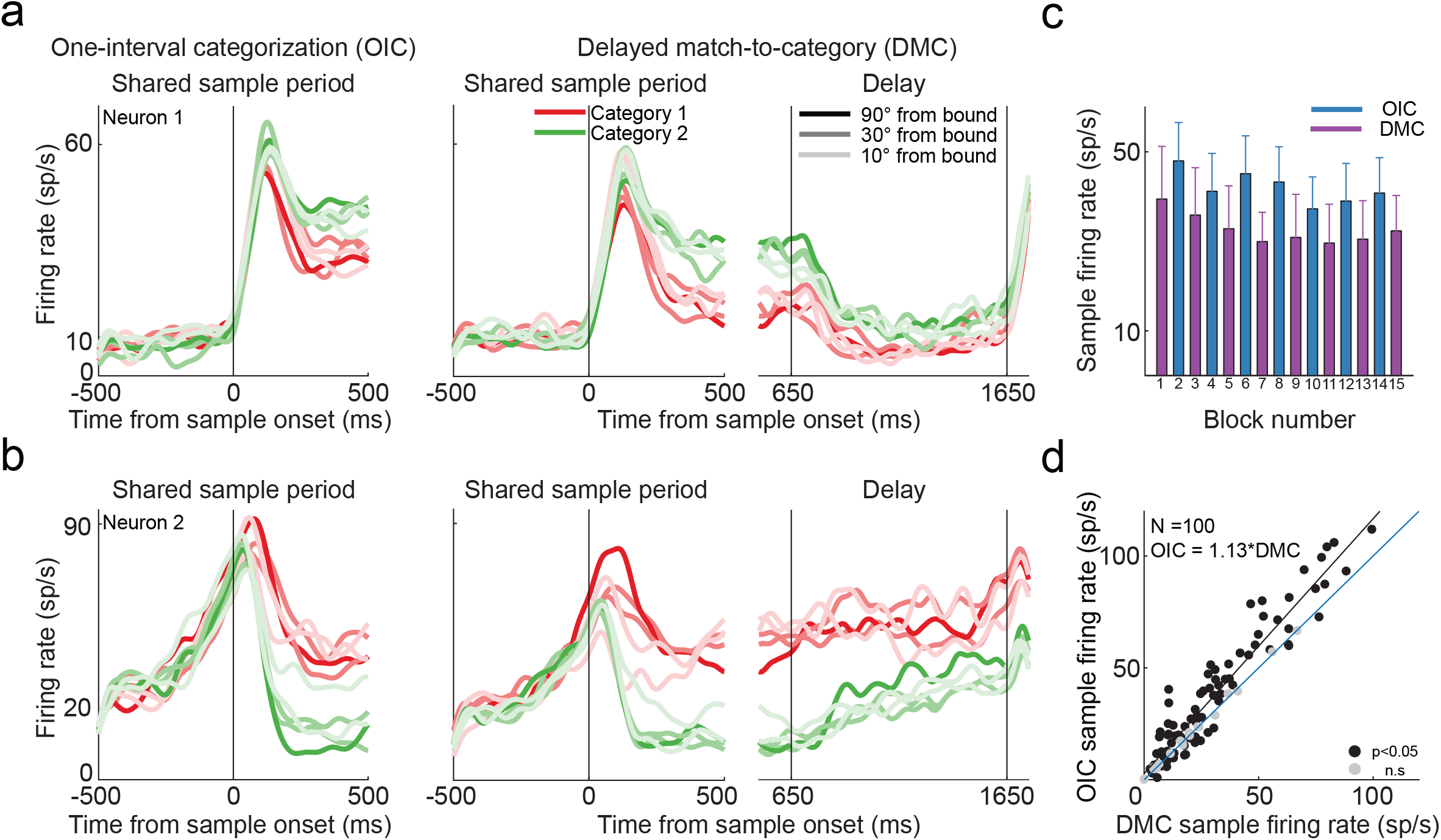
Single-neuron LIP responses in OIC and DMC. (a) Peri-stimulus time histogram (PSTH) of an example LIP neuron during the shared sample period in OIC (left) and DMC (center) and during the delay period in DMC (right). Spike trains were convolved with a Gaussian kernel (σ=30 ms). Colors represent categories and shade represents distance from the boundary, lighter shades refer to motion directions closer to the category boundary. Vertical lines at 0 indicate sample onset. Vertical lines at 650 and 1650 ms in the delay panel indicate delay onset and delay offset, respectively. (b) PSTH of another example neuron with plotting conventions as in 2a. (c) Block-wise comparison of firing rates during the shared sample period (0-500 ms from sample onset, offset by a visual latency of 80 ms) for the single neuron shown in 2a. Error bars represent s.e.m. (d) Scatter plot comparing OIC and DMC firing rates of the LIP population during the shared sample period. The black line is the regression line for all significant neurons (black dots, p<0.05, two-sample t-test; grey dots, not significant (n.s)), the blue line is the reference unity line with slope=1.

Higher average firing rates in the OIC than DMC task could result from multiple factors that vary between the two tasks, such as differences in motor response modality, task difficulty, task demands or expected value of reward. While we cannot isolate the factor(s) which contributes towards higher firing rates in OIC, we test one candidate. Because of LIP’s known involvement in saccadic eye movements, we hypothesized that LIP responses could be modulated by prior information about the motor modality used to report decision. Even before sample onset on each trial, the color of the fixation point indicated which task (and subsequently, effector) is relevant during that block. If LIP neurons were modulated by prior knowledge about the task or motor modality, we would expect fixation period firing rates to be higher in the OIC than DMC task. Out of 100 recorded neurons, 57 neurons showed significant differences in firing rates between the OIC and DMC tasks in the fixation period, of which 68.4% (39/57) showed greater activity in the OIC task (unpaired, two sample t-test, p<0.01). These observations suggest that LIP neurons could be preferentially engaged (i.e. have higher firing rates) when the task requires a saccadic (OIC) versus manual (DMC) report. It is possible that task difficulty influences firing rates such that differences in behavioral performance between the OIC and DMC tasks could explain firing rate differences. We controlled for task difficulty by considering a subset of task conditions, motion directions that were 30° away from the boundary, that were matched in accuracy and thus, were equally difficult in both tasks. For these accuracy matched conditions, we found that mean firing rates were significantly higher in the OIC task, suggesting that task difficulty cannot explain differences in the magnitude of firing rates between tasks (unpaired two sample t-test, p<0.01). However, as the OIC and DMC tasks differ in other aspects besides motor response modality, we cannot be certain which factors account for this activity difference. For instance, since OIC trials are shorter than DMC trials, monkeys received more reward per unit time during the OIC blocks than DMC blocks. This difference in reward frequency could have led to increased motivation in the OIC task which in turn could have led to increased firing rates.

### LIP neurons simultaneously encode multiple task-relevant variables

We next asked how encoding of task-relevant variables differed between tasks. We compared the fractions of cells that encoded the variables shared across tasks – sample motion direction and sample category. Since motion direction is nested within motion category as any given motion direction is associated with either category one or two but not both, we used a two-level nested ANOVA to quantify direction and category selectivity independently of each other. In the OIC task, 65% of recorded neurons were selective for motion direction (two-level nested ANOVA with direction nested within category, p<0.01) during the sample period while 50% were selective during the saccade epoch. Similarly, during the DMC task, 51% of recorded neurons were motion direction selective during the sample period and 25% were selective during the delay period. In addition to encoding motion direction, LIP neurons also encoded motion category throughout the trial. While 56% of cells were category selective during the OIC sample period, 63% of cells were category selective during the DMC sample period. In both tasks, many single neurons were category selective – responding similarly to directions within a category but distinctly to directions in different categories (example neurons in Figure 2a, 2b). Further, 40% of single neurons were direction selective in both tasks and 47% of neurons were category selective in both tasks. Finally, LIP neurons also encoded choice – i.e. saccade towards versus away from the RF in OIC and match versus non-match in DMC; 64% of neurons encoded saccade direction in OIC and 61% of neurons encoded match versus non-match in DMC. Thus, LIP activity multiplexes a variety of behaviorally relevant sensory (motion direction), decision (motion category), and motor (saccade direction and touch bar release) variables.

### Neural populations show stronger category representations in the DMC task

While single neurons encoded combinations of task-relevant variables in both tasks, we wondered whether population level category representations were also encoded similarly between tasks, since both tasks relied on the same category rules (i.e. category boundary). Specifically, we asked whether the strength and timing of category signals differed between the OIC and DMC tasks by quantifying category selectivity in two ways – (i) a category tuning index (CTI) applied to single neurons (Freedman and Assad, 2006) and (ii) linear decoders to classify motion category at the population level (Swaminathan, Masse and Freedman, 2013; Sarma *et al.*, 2015).

The CTI quantifies category selectivity for each neuron by comparing firing rates between pairs of motion directions within the same category (within category difference or WCD) versus different categories (between category difference or BCD). To measure the strength of category selectivity, we constructed the CTI by taking the difference between BCD and WCD and dividing by their sum. CTI values range from −1 to +1 with 1 indicating “binary-like” responses to categories (large differences in firing rate for directions in different categories) and −1 indicating zero differences between categories (large differences in firing rate for directions within each category). In both tasks, mean CTI values during the shared sample period were shifted towards positive values (indicating categorical tuning) and were significantly greater than values during the fixation-epoch (Figure 3a; OIC: sample CTI = 0.13 versus fixation CTI = 0.01; DMC: sample CTI = 0.19 versus fixation CTI = −3×10^−3^; paired t-test p<0.005 in both OIC and DMC).

**Figure 3.**
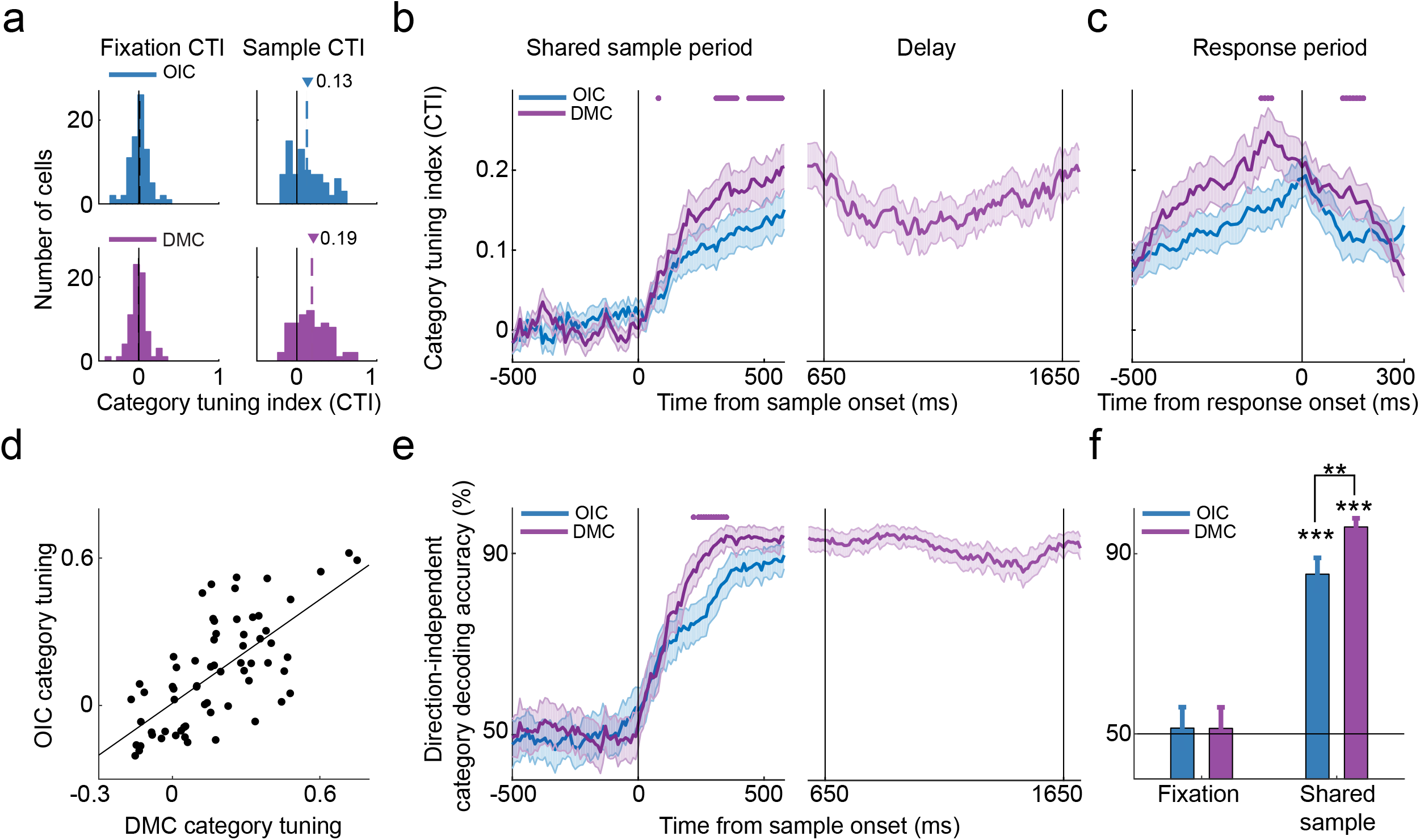
Stronger category tuning in DMC than OIC task. (a) Histograms of the category tuning index (CTI) during fixation and shared sample epochs in OIC and DMC tasks. The black solid line represents zero and the colored dashed lines represent mean CTI of direction selective neurons in each task (blue for OIC, purple for DMC). (b) Time-course of CTI computed by aligning firing rates to sample onset (left) during the shared sample period in OIC and DMC and the delay period (right) in DMC. (c) Time-course of CTI computed by aligning firing rates to response onset in OIC (saccade onset) and DMC (touch-bar release). In 2b and 2c, purple dots repre-sent timepoints at which traces are significantly different from each other with DMC > OIC (p<0.05, FDR-cor-rected) and shaded error bars indicate s.e.m. (d) Scatter plot comparing CTI in OIC and DMC for individual neurons during the shared sample period. The black line is the regression line indicating a significant, positive correlation between CTI values in both OIC and DMC tasks. (e) Time-course of performance of the direction-in-dependent category decoder trained and tested on OIC and DMC during the shared sample period (left) in OIC and DMC and the delay period (right) in DMC. Purple dots represent timepoints at which traces are significantly different from each other with DMC > OIC (p<0.05, bootstrap). Shaded error bars indicate s.d. (f) Mean performance of the direction-independent category decoder during fixation and shared sample epochs. Error bars indicate s.d. ** p<0.01, *** p<0.001, bootstrap.

Further, we examined the relationship between the strength of category selectivity in the OIC and DMC tasks. If individual neurons encoded category signals in one task, but not the other, we might find a negative correlation between CTI measures across the population. Instead, we found a positive correlation between CTI values of neurons in both tasks (Figure 3d; r^2^ = 0.68, p<10^−4^), indicating that the strength of category selectivity covaries between tasks.

Next, we compared the time course of category signals for all direction selective neurons in both tasks by calculating CTI in 200-ms windows stepped every 10 ms during fixation and shared sample periods. We found that mean CTI values were significantly greater in the DMC than OIC task during the late sample period, indicating stronger category representations in the DMC task (Figure 3b). Further, category selectivity was maintained through the delay period in the DMC task (Figure 3b – right panel). We also compared the time course of category signals when firing rates were aligned to response onset instead of sample onset, aligning neural responses to saccade onset in the OIC task and touch bar release in the DMC task (Figure 3c). We found that mean CTI values were greater in the DMC than OIC task even in the motor response-aligned condition. Interestingly, the time course of the response-aligned CTI monotonically increased to reach its maximum value just before movement onset in both OIC and DMC tasks, indicating that categorical tuning peaks immediately prior to the motor response, independent of the motor modality used to report the decision (eye versus hand).

We also used a population-level decoding approach to confirm our findings from the single-neuron CTI measure. We quantified the amount of category information in the LIP population using linear support vector machine (SVM) (Cortes and Vapnik, 1995) classifiers trained to decode whether population responses were elicited by stimuli from category one or category two. This analysis revealed strong and similar category decoding in both tasks (Figure S1, category decoder: DMC 99.4%, OIC 97.8%, bootstrap, p<0.001 compared to chance). However, this category decoder measured category selectivity without controlling for motion direction selectivity, an inherent property of LIP neurons, thus potentially overestimating the amount of explicit category information in the population. To reduce the influence of direction tuning on our measure of category selectivity, a different linear classifier was trained and tested on subsets of motion direction pairs with equivalent angular distances in each category. This direction-independent category decoder revealed higher category information in the DMC task as both the mean and time course of decoding accuracy were significantly greater in the DMC task (Figure 3e, 3f, category decoder: DMC 95.9%, OIC 85.5%, bootstrap, p<0.001 compared to chance). In sum, both the single-neuron CTI measure and the population-level decoders reveal that category selectivity is higher in the DMC task.

### Neural populations show stronger within-category direction representations in the OIC task

Greater category selectivity in the DMC task was unexpected because firing rates in the DMC task were lesser than in the OIC task. To understand which aspects of tuning accounted for greater category selectivity in the DMC than OIC task, we separately examined the BCD and WCD measures that were used to construct the category tuning index. This revealed that, while BCD values were comparable in both tasks, WCD values were significantly greater in the OIC than DMC task, indicating greater tuning for directions within a category in the OIC task (Figure 4a). Hence, the increase in CTI during the late sample of the DMC task is a direct consequence of more uniform firing rates among the within-category directions, i.e., within-category compression, resulting in a tendency for more binary-like category selectivity in the DMC task.

**Figure 4.**
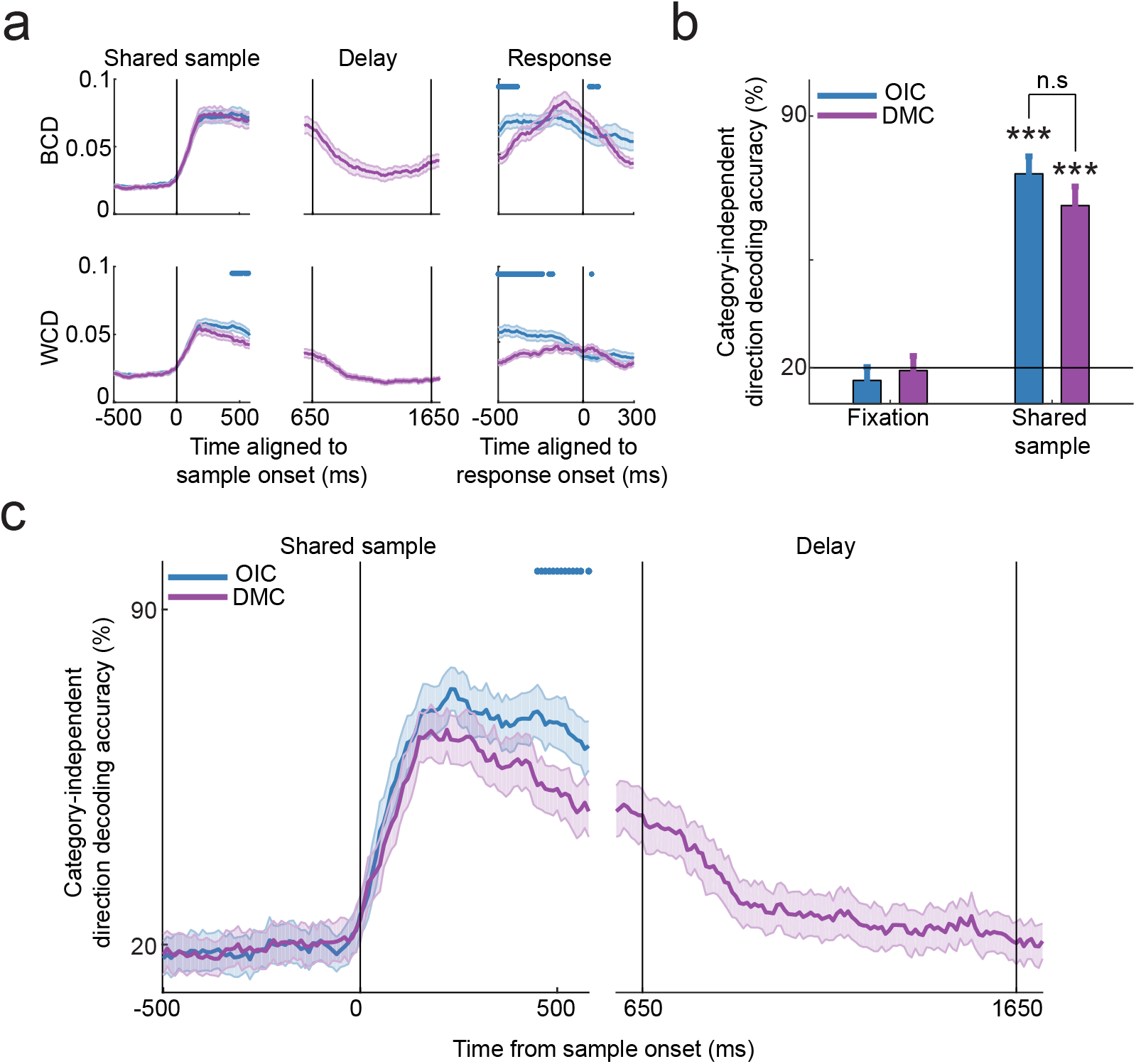
Stronger within-category direction tuning in OIC than DMC task. (a) Time-course of between-category difference (BCD, top row) and within-category difference (WCD, bottom row) computed by aligning firing rates to sample onset (left and middle column) and response onset (right column) in OIC and DMC (200 ms sliding window, stepped by 10 ms). Response onset is defined as saccade initiation towards the RF in OIC and touch-bar release in match trials in DMC. Vertical lines indicate sample onset in the left column, delay onset and delay offset in the middle column and response onset in the right column. Blue dots represent timepoints at which traces are significantly different from each other with OIC > DMC (p<0.05, FDR-corrected). Shaded error bars indicate s.e.m. (b) Mean performance of the category-independent direction decoder during fixation and shared sample epochs in OIC and DMC. Error bars indicate s.d. *** p<0.001, bootstrap. (c) Time-course of performance of the category-independent direction decoder trained and tested on OIC and DMC during the shared sample period (left) in OIC and DMC and the delay period (right) in DMC. OIC has more within-category direction information than DMC. Blue dots represent timepoints at which traces are significantly different from each other with OIC > DMC (p<0.05, bootstrap). Shaded error bars indicate s.d.

Greater WCD in the OIC task predicts that neural populations would encode more within-category direction information in the OIC than DMC task. Using category-independent direction decoders trained to classify motion directions within a category, we confirmed greater within-category direction information in the OIC task (Figure 4b, direction decoder: DMC 65.1%, OIC 73.9%, bootstrap, p<0.01 compared to chance). The time-course of direction decoding also showed higher accuracy for within-category direction information in the OIC than DMC task, particularly in the late sample period (400-500 ms from sample onset, Figure 4c). In contrast to the sustained category information present through the delay period in the DMC task, within-category direction information decreased during the same delay epoch, suggesting that behaviorally-relevant categorical information is stored in a low-dimensional, categorical representation instead of a high-dimensional, directional representation during the delay period (Figure 4a, 4c). Our results suggest that stronger category tuning in the DMC task is due to tighter clustering of responses to directions within each category that leads to larger differences between categories.

### Neural populations show more temporally stable category representations in the DMC task

Our results so far reveal stronger category representations in the DMC than OIC task. However, it remains unclear whether shared or distinct neural subpopulations encode category in the two tasks. In other words, are the same neurons contributing towards categorical judgments in a similar manner in both tasks? We employed a “cross-task” population decoding approach to directly compare the extent of overlap in population level representations in both tasks. We trained category decoders on population firing rates from one task (e.g. OIC) and tested the decoder on trials from the other task (e.g. DMC). The cross-task decoder reliably decoded category well above chance during the shared sample period in both tasks (Figure 5a, 5b, category cross-decoder testing on: DMC 77.6% (trained on OIC), OIC 81.4% (trained on DMC), bootstrap, p<0.01 compared to chance), indicating that the LIP population encoded categorical information in a similar manner using highly overlapping pools of neurons in the two tasks. In both tasks, the time-course of category decoding was nearly identical to the case in which the decoder was trained and tested on the same task (i.e. a “within-task” decoder) in the early sample period (0-250 ms from sample onset), but diverged in the late sample period (250-500 ms from sample onset). These findings suggest that category computations are supported by a common subpopulation of neurons in the early sample period and different subpopulations of neurons or different readout mechanisms in the late sample period.

**Figure 5.**
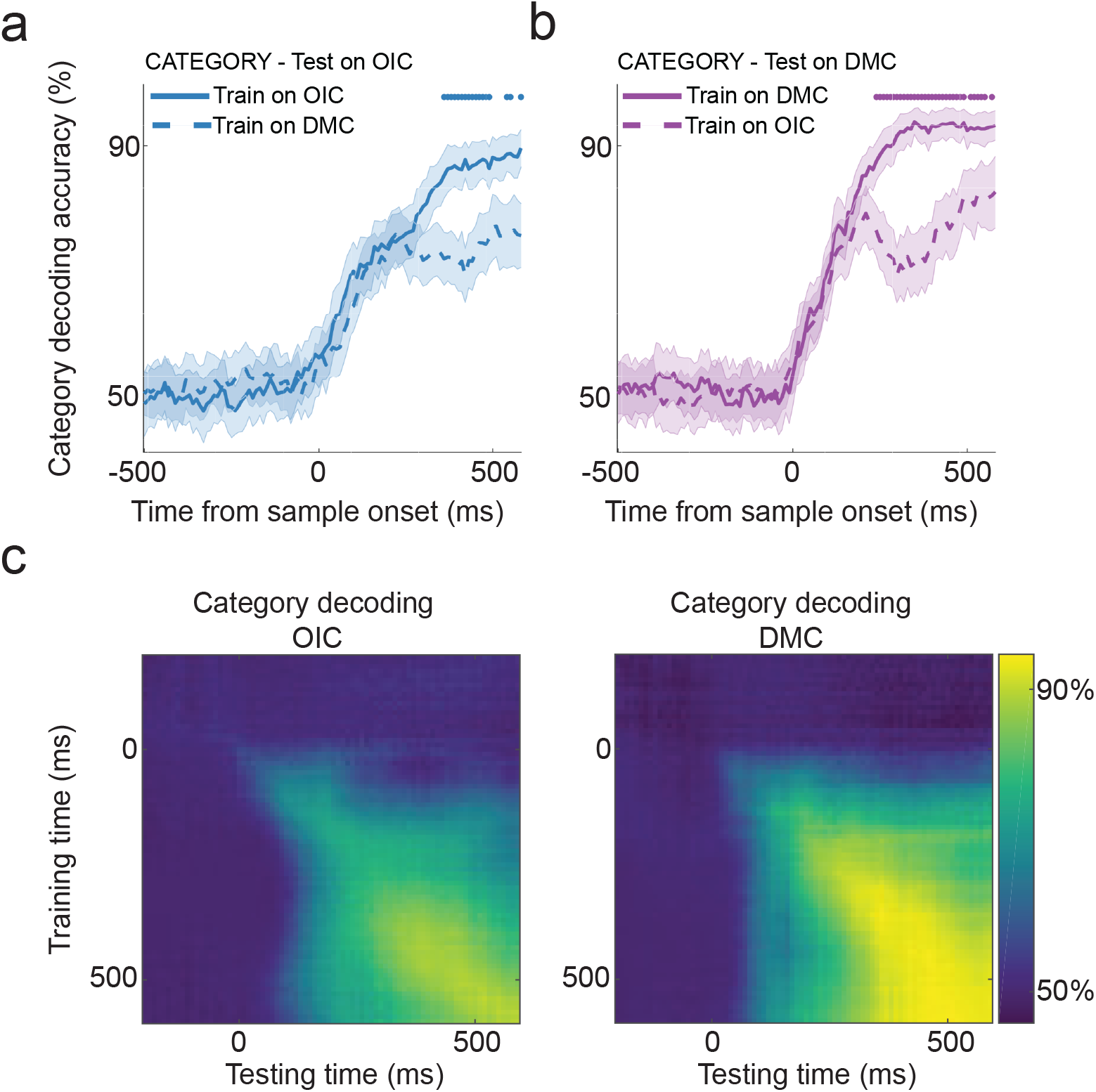
Temporal dynamics and stability of category decoding in OIC and DMC. (a-b) Time-course of performance of the category decoder tested on one task, but trained on data from the other task. In 5a, the category cross-decoder is trained on OIC or DMC, but always tested on OIC. In 5b, the category cross-decoder is trained on OIC or DMC, but always tested on DMC. Dots represent timepoints at which the cross-task decoder was significantly worse than the within-task decoder (p<0.05, bootstrap). Shaded lines represent s.d. (c) Performance of a cross-temporal category decoder trained and tested at all time points during fixation and shared sample epochs in OIC (left) and DMC (right). Coordinates (x,y) on the heatmap represent x=timepoint at which decoder was tested, y=timepoint at which decoder was trained.

We then sought to understand in more detail how representations differed in the late sample period. The late sample period is contextually distinct between the two tasks. In the OIC task, it is followed by a saccade for reporting sample category. In the DMC task, it is followed by an additional 150 ms of sample presentation, one second memory delay period, one or more test stimuli and a manual release of the touch bar. We wondered whether the differences in categorical encoding between the two tasks could be explained by the working memory demands during the delay period of the DMC task. Previous studies have found that during the delay period, neurons stably maintain their firing rates through persistent activity even after the stimulus is no longer present (Gnadt and Andersen, 1988; Funahashi, Bruce and Goldman-Rakic, 1989; Goldman-Rakic, 1995). Since firing rates remain stable over time during the delay period, encoded stimulus information at one time point can be used to infer stimulus information at a different time point during the delay, thus leading to temporally stable representations.

Consequently, we hypothesized that the requirement to maintain information during the delay in the DMC task would lead to persistent, temporally stable activity in anticipation of the upcoming delay period as the network gets ready to store the sample category. We therefore predicted that delay-dependent persistent activity in the DMC task would lead to greater temporal stability in the DMC task even before the delay, i.e. during the shared sample period. To ascertain whether categories are stored in more temporally stable format in the DMC task, we evaluated the temporal stability of category decoding using SVM decoders that were trained at one time point and then tested at all other times points in the shared sample period. The cross-temporal decoders revealed greater stability in the DMC than OIC task in the late sample epoch, as shown by higher decoding accuracies in the DMC task when decoders were trained and tested at different time points (Figure 5c).

These observations highlight two findings: first, neural activity and representations supporting sensory stimulus evaluation are similar between different tasks (OIC versus DMC) in the early sample period; second, neural population representations diverge following the initial evaluation of the sample stimulus, as the two tasks begin to differ in their behavioral demands. Notably, the population-level representations are governed by a stable, persistent categorical code in the DMC task, but a less temporally stable, more dynamic categorical code in the OIC task.

### RNN models recapitulate task-specific neural codes for category representations

Our analyses of LIP neural data showed greater category selectivity in the DMC task that was achieved by compressing variability among directions within a category. Further, category representations in the DMC task were characterized by greater temporal stability in the late sample period, thus resulting in a stable categorical code across time. In contrast, during the same shared sample period, OIC category representations exhibited more graded responses to directions within a category. Further, they were characterized by more transient encoding resulting in a more dynamic categorical code. What explains the differences between the underlying neural codes in the OIC and DMC tasks? We hypothesized that cognitive demands, specifically working memory demands, differentially modulate neural activity and dynamics to produce distinct neural codes. Since the sample category needs to be maintained in short-term memory during the delay period, we reasoned that the format of delay-period stimulus encoding would be affected by attractor dynamics which are thought to govern persistent activity generated by neural networks (Hopfield, 1982). The attractor dynamics pull neural activity into one of two states (corresponding to the two categories in the DMC task), which might reformat or compress category-related information to a simpler, binary format by collapsing all directions within a category to a single uniform category representation.

Inspired by recent approaches using artificial neural network models to understand task-related neural encoding and dynamics (Mante *et al.*, 2013; Chaisangmongkon *et al.*, 2017; Masse *et al.*, 2019), we tested whether recurrent neural networks trained to perform the OIC and DMC tasks reflected attractor-like dynamics that leads to compressed category representations in the DMC task. We trained 10 recurrent neural networks to solve the OIC task and 10 additional networks to solve the DMC task with the sequence of task events matched to the tasks used in the neural recording experiments (Figure 6a). These models received input from 36 motion-tuned neurons that projected to one recurrent layer consisting of 100 hidden neurons (80 excitatory + 20 inhibitory neurons). The excitatory neurons in the recurrent layer projected to three output units. The first output is a fixation neuron that was trained to remain active until response time which was 500 ms after sample onset in the OIC task and test stimulus onset in the DMC task. The other two output units in the OIC task reported category identity (category one or category two), while in the DMC task, they reported match or non-match status of test stimuli.

**Figure 6.**
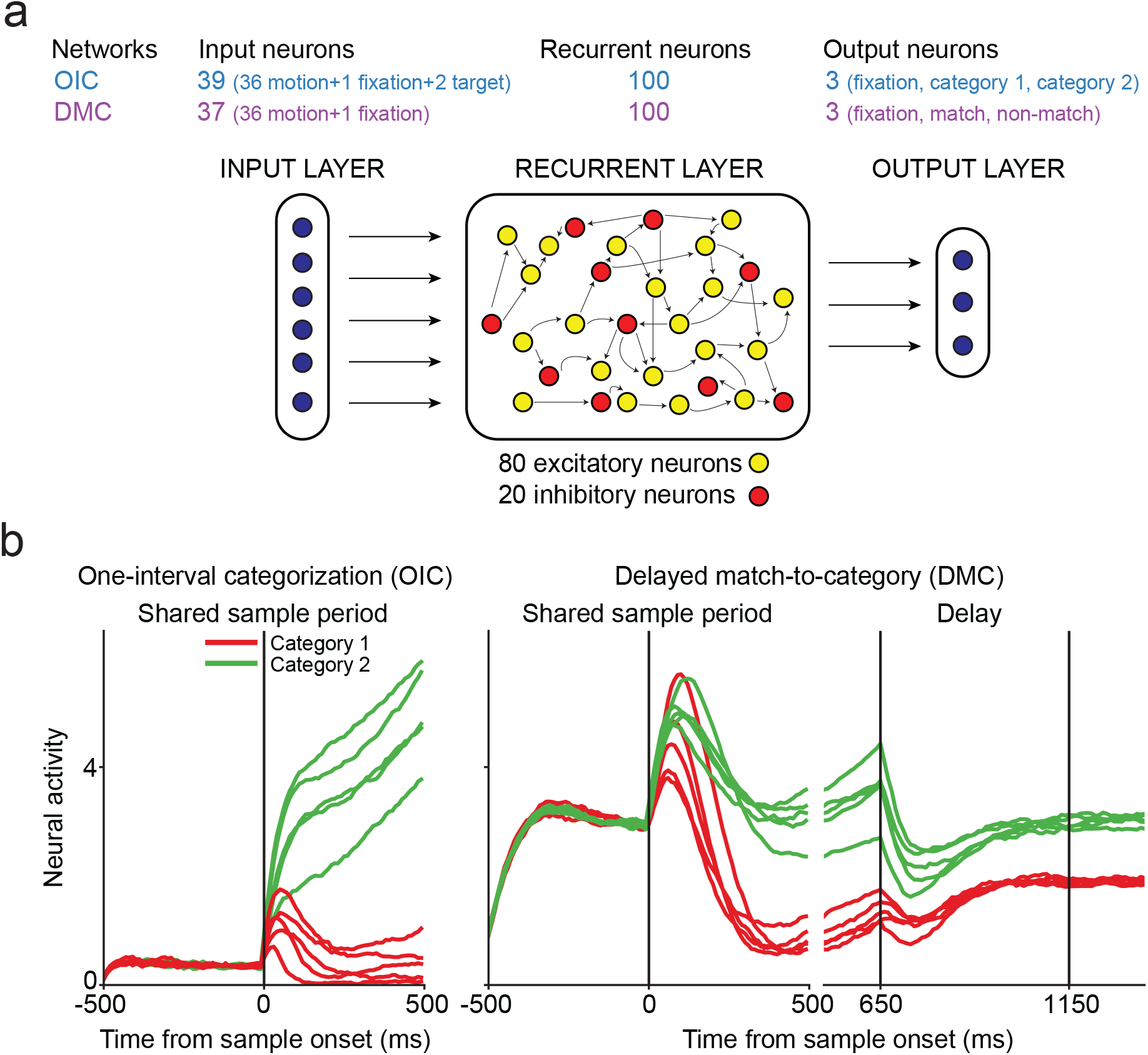
RNN model architecture and artificial unit responses in OIC and DMC. (a) Trained RNN models consisted of 36 motion-direction tuned input neurons which projected to 100 hidden neurons (80 excitatory+20 inhibitory) which in turn projected to three output units. In addition to the 36 motion-tuned units, OIC networks consisted of one fixation unit and two target units and DMC networks consisted of one fixation unit. In networks trained on OIC, output units corresponded to fixation, category one, and category two. In networks trained on DMC, output units corresponded to fixation, match, and non-match. (b) Peristimulus time histogram of two example RNN hidden units –one each from networks trained on OIC (left) and DMC (middle and right panel). Colors represent categories. Vertical lines at 0 indicate sample onset. Vertical lines at 650 and 1150 ms in the delay panel indicate delay onset and delay offset in DMC networks, respectively.

Trained networks successfully learned to optimize the desired output functions and the responses of the neurons in the hidden layer of these networks showed patterns of selectivity and dynamics which appeared similar to real neural data from LIP in these tasks. Applying the same category tuning analysis used for the LIP data on the artificial units from the RNN, we found OIC and DMC model neurons encoded category with similar responses to directions within a category and dissimilar responses to directions between categories (Figure 6b). We calculated individual units’ values of the category tuning index (CTI) which measured the degree to which units’ activity showed binary-like category selectivity. This revealed greater CTI values in networks trained on the DMC than OIC task, indicating more binary-like selectivity in the DMC task, akin to LIP neural populations (Figure 7a, 7c). Furthermore, we compared BCD and WCD values that were used to compute the CTI values and found that BCD values were similar in networks trained on both tasks, but WCD values were significantly greater in the networks trained on the OIC than DMC task, suggesting greater compression of activity among direction within categories in the DMC task (Figure 7b). Thus, stronger binary selectivity could result from greater within-category compression, as a result of attractor dynamics underlying short-term memory needed to bridge the delay period in the DMC task. This suggests that differences in memory-dependent cognitive demands between tasks could produce task-specific differences in the format of underlying neural codes.

**Figure 7.**
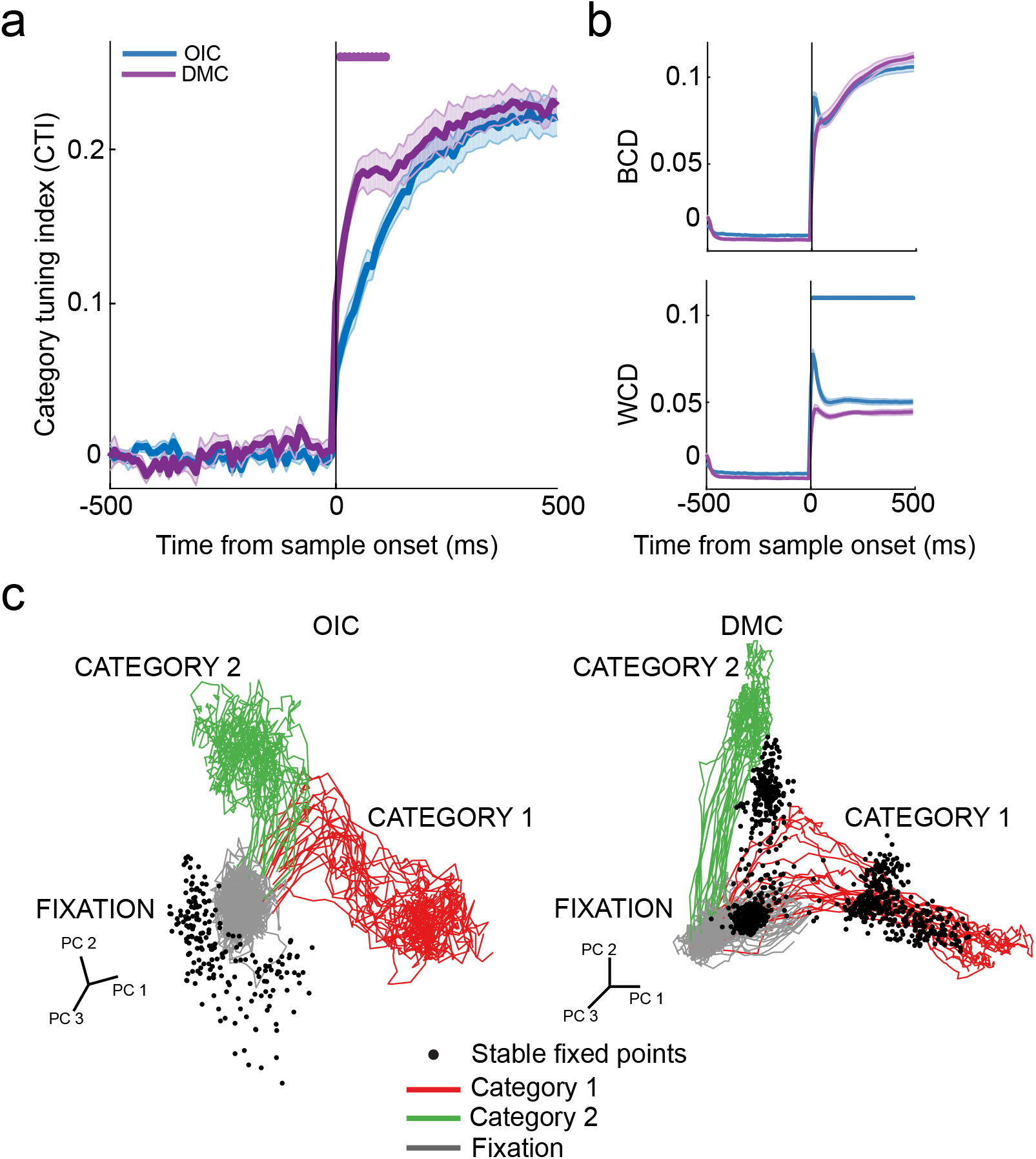
Categorical encoding and dynamics in RNNs trained on OIC and DMC tasks. (a) Mean time-course of the category tuning index (CTI) during the shared sample period averaged over 10 RNNs trained independently on OIC and DMC. Purple dots represent timepoints at which traces are significantly different from each other with DMC > OIC (p<0.05, FDR-corrected). (b) Mean time-course of between-category difference (BCD, top row) and within-category difference (WCD, bottom row) during the shared sample period averaged over the same 10 RNNs shown in 7a. Blue dots represent timepoints at which traces are significantly different from each other with OIC > DMC (p<0.05, FDR-corrected). In 7a and 7b, vertical line at 0 indicates sample onset and shaded error bars indicate s.e.m. calculated over 10 networks. (c) Fixed-point structure overlaid with hidden-layer neural trajectories of an example RNN model trained on OIC (left) and DMC (right) tasks separately. Colored traces indicate PCA trajectories of the hidden layer neurons during fixation and shared sample epochs. Each trace corresponds to a single trial with fixation colored in gray, category one trials in red and category two trials in green. Black dots indicate stable fixed points of the trained networks computed on the hidden layer neural activity during the shared sample period.

To understand whether different neural codes reflect different computational mechanisms, we explored model dynamics through the analysis of fixed points in state space. Fixed points (small black dots, Figure 7c) correspond to neural activity patterns that are stable when external sensory inputs are turned off. A recent study performed fixed point analysis on RNN models trained on the DMC task and found that the dynamical mechanisms consisted of stable states at the end of the sample period, followed by a dynamic, high-velocity trajectory during the delay period (Chaisangmongkon *et al.*, 2017). Similarly, in the absence of external input, we also found stable states associated with each category at the end of the sample period in the DMC task. In addition, we also found a stable state in the fixation period. This dynamical analysis revealed that in a delay-based categorization task, stable states that may reflect attractors emerge as sample categories need to be maintained in working memory. Indeed, at the end of the sample period, population activity for all motion directions within a category converges towards the corresponding stable category state, resulting in compression of stimulus representations in anticipation of the upcoming delay period. In contrast, category-based stable states were not present in networks trained to perform the OIC task in which there are no explicit working memory demands. Instead, in the absence of input, there was one stable state that coincided with the fixation period. These results suggest that attractor dynamics underlying working memory could contribute to stronger within-category compression in the DMC task.

## Discussion

We examined the neural correlates of abstract categories during flexible task switching between two categorization tasks which varied in their behavioral demands. Both tasks required monkeys to categorize visual motion stimuli according to the same category boundary. However, the tasks varied in whether category judgments were reported immediately (OIC) or whether they were based on sequential comparisons of sample and test categories (DMC). The tasks also varied in their motor response mappings, demands on short-term memory, and the effector (eye or hand movement) used for the decision report. Despite these differences in task demands, the categorization requirements of each task during the sample presentation are similar. We examined whether LIP plays a general role across both tasks, or whether LIP is more specialized for task-specific demands, such as working memory, sequential comparison, or motor response modality.

We found that LIP activity robustly encodes motion categories using a common coding scheme in both tasks with stronger category representations evident in the DMC task. In the DMC task, neurons exhibited stronger within-category compression by responding with more homogeneous firing rates for directions within each category. By contrast, neurons in the OIC task responded with more graded patterns of activity for directions within each category. This produced more binary-like or abstract category representations in the DMC task, and more mixed encoding of direction and category in the OIC task. Furthermore, during the shared sample period in the DMC task, categorical encoding was more temporally stable and sustained, presumably reformatting category encoding in preparation for maintaining information in the imminent delay period. In contrast, categorical encoding during the OIC task was more transient, presumably as sensory and decision information is rapidly routed to motor circuits in that task. In sum, our results suggest that LIP circuits contribute to flexible task switching through generalized category representations that are rapidly reconfigured and specialized to meet task demands.

Previous studies in LIP have focused on perceptual decisions, often employing behavioral paradigms with relatively simple stimulus-response (S-R) mappings in which noisy sensory stimuli are mapped directly to motor responses (Gold and Shadlen, 2007; Antzoulatos and Miller, 2011). Recent studies have also investigated more flexible mappings between stimuli and motor responses (Gold and Shadlen, 2003; Bennur and Gold, 2011). Other studies have employed multiple response modalities to report the same decisions (de Lafuente, Jazayeri and Shadlen, 2015) or modulated behavioral demands in both cued (Mante *et al.*, 2013; Siegel, Buschman and Miller, 2015; Kumano, Suda and Uka, 2016) and un-cued (Snyder, Batista and Andersen, 1997; Asaad, Rainer and Miller, 1998) scenarios. Flexible decisions have also been studied in PFC and PPC by training animals to alternate between tasks with different sensory stimuli, different decisions and different motor responses in order to understand how circuits route information according to shifting behavioral demands. In both the OIC and DMC tasks in our study, the category decision is decoupled from the eventual motor action, allowing us to delineate LIP’s role in decision-making and response selection. In contrast to earlier work finding no impact of LIP inactivation on a motion discrimination task during simple S-R decisions (Katz *et al.*, 2016), a recent study found that inactivation of LIP neurons indeed impairs perceptual decisions in a task with flexible S-R mappings (Zhou and Freedman, 2019). Although incorporating cognitive flexibility in tasks might engage new and task-specific neural mechanisms that might not generalize to simpler tasks, this study suggests that introducing flexibility in classical task structures can provide a more nuanced understanding of decision-making circuits.

Abstract, categorical signals have been observed in multiple brain regions including PFC (Freedman *et al.*, 2001; Swaminathan and Freedman, 2012), PPC (Freedman and Assad, 2006), and IT cortex (Freedman *et al.*, 2003). Recent studies found that both PFC and ITC show category-correlated encoding during a shape DMC task; however, PFC neurons encode shape categories in a more abstract, binary format whereas ITC encoding was in a more mixed format in which categorical information was mixed with visual feature information (Meyers *et al.*, 2008). These results bear a striking resemblance to the task-specific coding schemes we find in this study, with LIP encoding revealing abstract, binary category representations in DMC and mixed, graded category representations in OIC. While these previous studies suggest that different brain regions might support distinct formats of categorical encoding, our results demonstrate that behavioral task demands can also produce different encoding formats within a single brain region. Previous categorization studies have proposed that the abstract category signals observed in the DMC task are the output of a neural computation that transforms sensory representations into explicit, binary category encoding, which explicitly represents the learned stimulus categories (Freedman and Assad, 2006, 2016). However, the results from this study present an alternative interpretation that the specific format of category signals – abstract and binary – observed in the DMC task may be a consequence of – (i) short-term memory demands of delayed matching tasks, and (ii) the dynamics of neural population activity in brain regions that support working memory.

Several studies finding abstract categorial encoding have also employed behavioral tasks that have a working memory requirement. One study found that, in addition to motion categories, LIP also encodes learned shape associations in an abstract format in a delayed pair association task (Fitzgerald, Freedman and Assad, 2011). Interestingly, the neurons showing strong delay-period encoding of motion categories in that task also often showed strong delay-period shape-pair encoding. PFC neurons also show abstract categorical encoding in a shape DMC task with three categories (as opposed to two), suggesting that abstract category signals can also extend to multiple groups or attractor states (Freedman *et al.*, 2002). Another study found binary-like encoding of task rules as rule-cue stimuli (both auditory and visual) evoked similar patterns of neural activity when they were associated with the same rule (Wallis, Anderson and Miller, 2001). Recent work also examined learned associative representations in LIP and found that across multiple tasks with a memory delay period, nearly all recorded neurons had a similar order of preference among associated stimuli, thus creating biased representations (Fitzgerald *et al.*, 2013). Such biased neural representations are predicted from a recent recurrent neural network model of LIP developed to explain the origins of persistent memory delay activity (Ganguli *et al.*, 2008). The model proposes that over long timescales, local LIP activity is one-dimensional such that the network dynamics relax to a single firing-rate mode as the firing rate stabilizes after a transient input, such as during a memory-delay period. Our results also suggest that such network dynamics compress directions within a category to a single point-attractor in the DMC task, but not OIC task. Working memory demands also restructure neural encoding and dynamics in the motor system as neural populations in the motor cortex achieve a low-dimensional preparatory state during delayed reaches, but bypass this step during non-delayed reaches (Ames, Ryu and Shenoy, 2014). Collectively, these observations suggest that the requirement to maintain task-relevant information in working memory could reformat stimulus representations, specifically through within-category compression to a single neural state or attractor.

Our study indicates that both abstract and mixed category encoding schemes coexist within LIP in a task-dependent manner with a role for recurrent dynamics in compressing neural categorical encoding to appear more abstract and binary. It is likely that this phenomenon is mediated by interactions among different brain regions involved in the OIC and DMC tasks.

Indeed, LIP is connected with the dorsolateral prefrontal cortex (DLPFC) (Chafee and Goldman-Rakic, 1998), frontal eye fields (FEF), and superior colliculus (SC) (Blatt, Andersen and Stoner, 1990; Lewis and Van Essen, 2000), all of which exhibit persistent activity during memory-delay periods. Furthermore, in concert with FEF and SC, LIP plays a significant role in oculomotor planning and spatial attention, and we predict that FEF and SC neurons would also be selective during target selection, saccade planning and initiation in the OIC task. Future studies with recordings in LIP, FEF, SC, and PFC will elucidate the roles of these different nodes in categorical decision-making, working memory and response selection.

This study examines how flexible cognitive demands such as working memory interact with neural encoding and dynamics to produce task-specific neuronal representations. We propose that attractor dynamics underlying short-term working memory reorganizes graded stimulus representations into binary-like categorical encoding during delayed matching tasks. This advances our understanding of the interactions between neural feature selectivity and dynamics, and how cognitive demands flexibly affect these dynamics in order to support flexible task performance.

## Acknowledgements

We thank S. Swaminathan for expert behavioral and experimental assistance and the staff at the University of Chicago Animal Resources Center for expert veterinary assistance. We also thank N. Masse, G. Ibos, W. Johnston, K. Latimer, Y. Zhou, B. Peysakhovich, M. Rosen, and P. Moghimi for their comments on an earlier version of this manuscript.

## Competing Financial Interests

The authors declare no competing financial interests.

## Methods

### Behavioral task and stimulus display

Two male monkeys (*Macaca mulatta*) were trained to alternate between a one-interval categorization (OIC) task and a delayed match-to-category (DMC) task in blocks of 20 trials (monkey B) or 30 trials (monkey M) based on a colored cue presented at the start of a trial. In both tasks, monkeys were trained to categorize the same random-dot motion stimulus into one of two categories. Stimuli were circular patches (~5° diameter) of ~100 high-contrast dots that moved with 100% coherence at a speed of 12°/s. Motion stimuli were created by dividing 360° of motion directions into two categories based on an arbitrary learned category boundary (Figure 1b). Six evenly spaced motion directions (60° apart; 15°, 75°, 135°, 195°, 255°, 315°) were used as sample and test stimuli in addition to four directions that were 10° away (35°, 55°, 215°, 235°) from the category boundary. In OIC, monkeys indicated the category membership of a sample stimulus by making an eye movement to a colored target associated with the category. In DMC, monkeys indicated whether sequentially presented sample and test stimuli matched in category by releasing or holding a manual touch-bar for match or non-match trials respectively.

In the OIC block, every trial started with a yellow colored fixation point (0.12° radius) at the center of the screen. After gaze fixation was maintained for 500 ms (within a 2°-radius fixation window), a sample stimulus was presented for 500 ms in the receptive field (RF) of the recorded neuron. At 500 ms after sample onset, two colored targets (red and green squares) appeared – one in the RF of the recorded neuron and the other 180° opposite to the RF. Once the targets appeared, the monkeys made a saccade and maintained fixation for 300 ms on the red target or the green target to indicate the category of the sample stimulus as category 1 or category 2 respectively, in order to receive a juice reward. Critically, on every trial, the red and green saccade target locations were counterbalanced between the two possible locations (inside RF or outside RF). This ensured that the saccade directions were not correlated with the category of the sample stimulus, thus, allowing us to disambiguate sensory and/or decision signals from motor signals in the sample period.

In the DMC block, every trial started with a white colored fixation point (0.12° radius) at the center of the display. After gaze fixation and touch-bar press was maintained for 500 ms (within a 2°-radius fixation window), a sample stimulus was presented for 650 ms in the receptive field of the recorded neuron, followed by a 1000 ms delay and a test stimulus presented for 650 ms. If the sample stimulus and the test stimulus matched in category, the monkeys released a manual touch-bar using a hand movement, in order to receive a juice reward. If the sample and the test stimulus did not match in category, a second test stimulus appeared, which was always a category match to the sample, and the monkeys were required to release the touch-bar.

Gaze positions were measured using an Eyelink 1000 optical eye tracker (SR Research) at a sampling rate of 1 KHz and stored for offline analysis. Task events, stimulus display, timings and reward delivery were controlled via a MATLAB-based toolbox, MonkeyLogic. Stimuli were displayed on a 21-inch color CRT monitor (1280×1024 resolution, 75 Hz refresh rate, 57 cm viewing distance).

### Electrophysiological recoding

Two male monkeys were implanted with a headpost and a recording chamber. The recording chamber was implanted over the intraparietal sulcus centered ~3 mm posterior to the inter-aural line. Stereotaxic coordinates were determined from anatomical MRI scans obtained prior to headpost and chamber implantation. All surgical procedures were in accordance with the University of Chicago’s Animal Care and Use Committee and US National Institutes of Health guidelines.

LIP recordings were conducted using 75-μm tungsten microelectrodes (FHC), a dura-piercing 27 Ga guide tube and an electronic micromanipulator (NAN Instruments). Neurophysiological signals were amplified, digitized, and stored for offline spike sorting (Plexon) to verify the quality and stability of neuronal isolation within a recording session.

### RF mapping and stimulus placement

In every recording session, we tested and recorded the activity of well-isolated neurons during the delayed memory-guided saccade (see below). We identified the lateral intraparietal cortex (LIP) based on the presence of neurons that showed spatially selective, delay, and/or peri-saccadic activity during the delayed memory-guided saccade task. The spatial location on the screen that elicited the highest activity was identified as the receptive field (RF). Stimuli in the OIC and DMC tasks were always presented inside the RF of the recorded neuron(s). The eccentricity of stimulus placement for LIP recordings ranged from 5-9°. The depth of LIP recordings ranged from 4-11 mm from the surface of the dura.

### Delayed memory-guided saccade task

In this task, every trial started with a white colored fixation point (0.12° radius) at the center of the display. After gaze fixation was maintained for 500 ms (within a 2°-radius fixation window), a white square was presented for 300 ms at a fixed eccentricity in one of eight possible peripheral locations. The brief target flash was followed by a 1000 ms delay after which the fixation point disappeared, cueing the monkey to make a saccade to the remembered location.

### Data analysis

All analyses were conducted on correct trials, excluding trials with incorrect responses, fixation breaks, and early responses. Unless otherwise stated, all our results were qualitatively and quantitatively similar in both monkeys. Thus, we combined datasets from both monkeys for all analyses.

### Epoch-based analysis

The fixation period was a 500-ms epoch prior to sample onset in both the OIC and DMC tasks. The sample period was a 500-ms epoch in the OIC task and a 650-ms epoch in the DMC task, both epochs offset by 80 ms to account for neuronal response latency. Since the length of the sample period was different in the two tasks, we analyzed all comparisons between the OIC and DMC tasks during the shared sample period which was a 500-ms epoch after the sample stimulus onset in both tasks. Early and late sample periods were defined as 250-ms epochs beginning at 80 ms and at 330 ms after stimulus onset. In the OIC task, the saccade/choice period was a 250-ms epoch starting at saccade target onset which was 500 ms after sample stimulus onset. In the DMC task, the delay period was a 1000-ms epoch beginning at sample stimulus offset and the match/non-match or choice period was a 250-ms epoch starting at test stimulus onset.

### Category tuning index (CTI)

We quantified category selectivity for individual neurons through a category tuning index that compared neuronal responses for motions directions in the same versus different categories. The CTI index was computed as the difference between two quantities – the between category difference (BCD) and the within category difference (WCD). BCD was measured as the average firing rate difference between pairs of motion directions in different categories. Similarly, WCD was measured as the average firing rate difference between pairs of motion directions within the same category. To account for the inherent motion direction tuning of LIP neurons, we considered only pairs of directions for which the angular difference was equal in BCD and WCD. Out of 25 (5×5 directions per category) possible direction pairs for BCD and 20 (2 categories × ^5^C_2_) possible direction pairs for WCD, the angular difference of 12 motion direction pairs were equal in BCD and WCD. CTI values ranged between −1 (directions within a category are much more distinct from each other compared to directions between categories) to +1 (directions in different categories are much more distinct from each other compared to directions within categories).

### Population decoding

We constructed linear population decoders using support-vector machine classifiers that were trained and tested to classify task-relevant variables including motion direction and motion category. Because the neural data were collected through single-electrode recordings and any individual recording session had 1-3 units, we concatenated units across sessions to create a larger pseudo-population of 100 neurons. In all the population decoding analyses, we used a four-fold cross-validation approach in which 75% of the data was used for training and the remainder for testing. The detailed methods used in the decoding analysis have been described previously and are briefly explained in the following sections(Sarma *et al.*, 2015).

#### Category-independent direction decoder

We measured direction information independently of category by constructing separate direction decoders for each category and averaging performance across both decoders. We randomly sampled with replacement 20 trials for each direction in a category and used an SVM multi-class classifier(Chang and Lin, 2011) (one versus one) to classify one among five directions within a category (chance performance is 20%).

#### Direction-independent category decoder

We measured category information independently of direction by constructing category decoders in which training and testing sets consisted of motion directions with equal angular distance from the boundary in both categories. We constructed different classifiers for each of the two motion directions within a category that were 30° and 10° away from the boundary and their diametrically (and hence categorically) opposite sample directions. These classifiers were trained on trials from the remaining sample directions in each category which have the same angular distance from the directions in the testing set, and then tested on the sets of opposing directions in the test set. For each motion direction, we randomly sampled 20 trials with replacement and trained the classifier to classify test response vectors as responses for category 1 or category 2 (chance performance = 50%). For the cross-task category decoding analysis, we constructed category decoders that were trained on trials from one task (e.g., OIC) and tested on trials from the other task (e.g., DMC).

For both the direction and category decoders, we repeated the resampling procedure 10,000 times for each epoch in all epoch-based analyses and 100 times for each time point in all sliding window analyses.

### Temporal stability analysis

We extended the category decoding approach to compare the amount of category information encoded at different time points in the trial using a 200-ms sliding window with a 20-ms step size. To assess if category information is maintained in population activity over time, we trained a direction-independent category decoder at one time point in the trial and tested it at all time points during the shared sample period. If the category decoders were able to decode category information at different time points during the shared sample period, this indicated that the patterns of neural activity were similar over time. A block-like structure of high decoding accuracy indicates stable patterns of category selectivity over the time-course of a trial.

### Network models

The details of the recurrent neural network modeling used in this study have been described previously and are described briefly in the following section(Masse *et al.*, 2019).

Neural networks were trained and simulated using the machine learning library TensorFlow(Abadi *et al.*, 2016). In both tasks, the inputs were visual motion stimuli moving in one of 10 equally spaced directions. Networks trained on both the OIC and DMC tasks consisted of 36 motion-direction tuned input neurons. Networks trained on the OIC task alone also contained inputs of two target units that specified the location of response targets (left/right) and one fixation neuron that was active until 500 ms from after sample onset. Networks trained on the DMC task also contained one fixation neuron that remained active until the end of the delay. These input neurons projected to a hidden layer consisting of 100 recurrently connected neurons (80 excitatory + 20 inhibitory). Networks trained on the OIC task contained three output neurons – one that represented the decision to maintain fixation, and two neurons that represented the selection of category 1/category 2. Networks trained on the DMC task also contained three output neurons – one neuron represented fixation, and two neurons represented match/non-match responses. Ten separate networks, in which the parameters of each network were randomly initialized, were trained on OIC and DMC tasks, with trial epochs that matched those performed by behaving animals. In order to make training easier, the delay period for the simulated DMC task was 500 ms, instead of the 1000 ms delay used in the neurophysiological experiments.

The neural activity in the hidden layer were governed by the following equation:

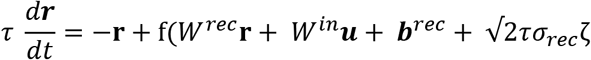

where τ is the neuron’s time constant, f(.) is the activation function, *W*^*rec*^ and *W*^*in*^ are the synaptic weights between recurrent neurons, and between input and recurrent neurons, respectively, ***b***^*rec*^ is a bias term, ζ is independent Gaussian white noise with zero mean and unit variance applied to all recurrent neurons and σ_*rec*_ is the strength of the noise. The differential equation was then discretized using a first-order Euler approximation with time step of 10 ms.

The networks were trained with the following parameters which were kept constant for both tasks: neuron time constant: 100 ms, time step: 10 ms, standard deviation of input noise: 0.1, standard deviation of recurrent noise: 0.5. gradient batch size: 256, number of batches: 2000.

Initial connection weights from the input layer, projecting to the output layer, and between excitatory neurons were randomly sampled from a gamma distribution with shape parameter 0.1 and scale parameter of 1. Initial connection weights projecting to or from inhibitory neurons were sampled from a gamma distribution with shape parameter 0.2 and scale parameter of 1. Initial bias values were set to 0. The networks were trained to minimize the cross-entropy between the actual and target output with Adam gradient descent optimization, and the mean L2 norm of the recurrent neurons’ activity level. All trained networks had accuracy greater than 95%.

Fixed-point analysis: For finding fixed points in the trained networks, we implemented FixedPointFinder(Golub and Sussillo, 2018), an open-source Tensorflow toolbox for finding fixed points from linearized dynamics in high-dimensional trained RNNs. Recurrent activity during the sample period of trained RNNs was used to find fixed points. We computed fixed points separately for OIC and DMC networks. They were calculated for 1024 initial points that were randomly sampled across trials, neurons, and time within a trial during the shared sample period. Joint optimization was utilized to find all fixed points simultaneously for all initial points. Fixed points computed during the shared sample period were visualized by overlaying then over trajectories of trials in PCA space. PCA was performed on the recurrent layer.

### Statistics

Selectivity for various task-relevant variables including motion direction, motion category was computed using a two-level nested ANOVA since direction is nested in category (p<0.05). Saccade direction selectivity was measured using a one-way ANOVA (2 saccade directions, into RF versus away from RF, p<0.05) and match/non-match selectivity was also measured using a one-way ANOVA (match versus non-match, p<0.05). Comparisons between proportions of direction selective or category selective neurons were conducted using a chi-square test.

The statistical significance of differences in CTI, BCD, and WCD was computed through a shuffle analysis. To obtain a null distribution, we shuffled the direction labels, calculated CTI and repeated this process 1000 times for both the OIC and DMC tasks. An CTI value was significantly different from 0 if it was greater than 95% of values from the null distribution. The statistical significance of differences in the time-course of CTI, BCD, and WCD was computed using a two-sided Wilcoxon rank sum test and we corrected for multiple comparisons using false discovery rate.

The statistical significance of differences in decoding accuracy for both direction and category information in OIC and DMC were determined using a bootstrap analysis. For the epoch-based decoding analysis, we constructed a null distribution by shuffling the labels for motion direction and then calculated decoding accuracy for the shuffled distribution. Average classification accuracy was considered significantly greater than chance if the value was greater than 95% (p<0.05) or 99% (p<0.01) of the values from the null distribution. For the time-course based decoding analysis, we used a bootstrap analysis to compare classification accuracies between OIC and DMC at each time point. Average classification accuracy was significantly different between OIC and DMC if 97.5% (p<0.05) or 99.5% (p<0.01) of values from one task were greater than the other task.

---

**Figure S1.**
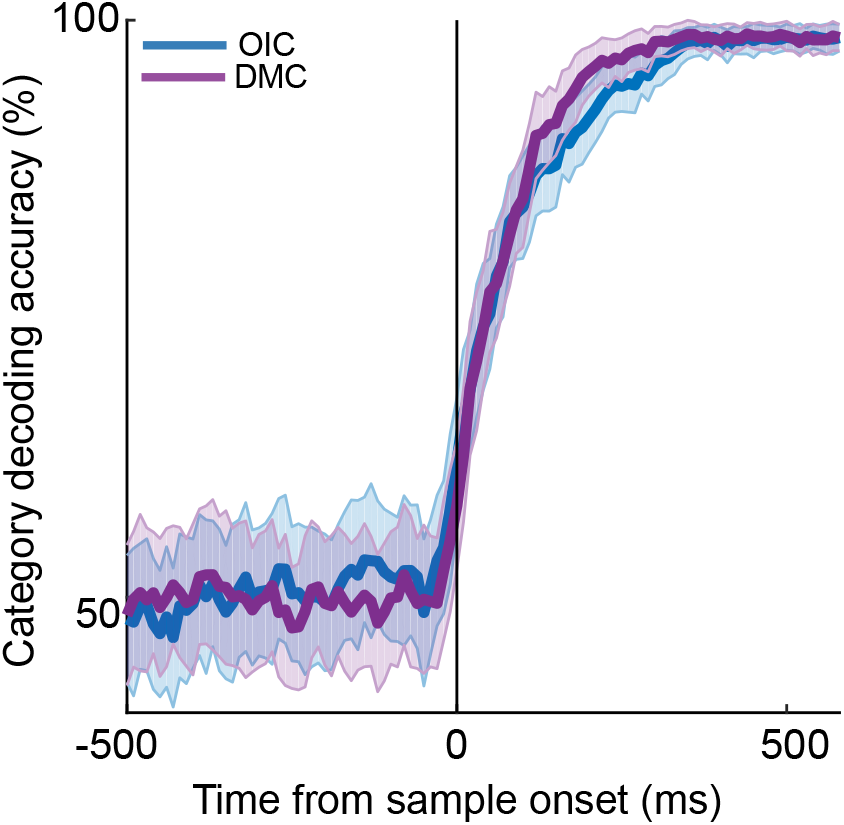
Strong and similar category tuning in LIP in both OIC and DMC. Time course of the performance of a category decoder trained and tested on OIC and DMC during the shared sample period. Shaded error bars indicate s.d.

## Notes

### Competing Interest Statement

The authors have declared no competing interest.

